# A method for stabilising the XX karyotype in female mESC cultures

**DOI:** 10.1101/2022.04.07.487567

**Authors:** Andrew Keniry, Natasha Jansz, Peter F. Hickey, Kelsey A. Breslin, Megan Iminitoff, Tamara Beck, Matthew E. Ritchie, Marnie E. Blewitt

## Abstract

Female mouse embryonic stem cells (mESCs) present differently to male mESCs in several fundamental ways, however complications with their *in vitro* culture have resulted in an underrepresentation of female mESCs in the literature. Recent studies show that the second X chromosome in female, and more specifically the transcriptional activity from both of these chromosomes due to absent X chromosome inactivation, sets female mESCs apart from males. To avoid this undesirable state, female mESCs in culture preferentially adopt an XO karyotype, with this adaption leading to loss of their unique properties in favour of a state near indistinguishable from male. If female pluripotency is to be studied effectively in this system, it is critical that high quality cultures of XX mESCs are available. Here we report a method for better maintaining XX female mESCs in culture, which also stabilises the male karyotype, and makes study of female specific pluripotency more feasible.

## Introduction

Female pluripotent stem cells differ from males genetically, epigenetically and functionally^1-8^. However, due to complications with *in vitro* culture of female mouse embryonic stem cells (mESCs), the vast majority of mESC research has been performed on male lines. The first confirmed mESC line was male^9^. Subsequently, lines employed as workhorse cells in the field, E14, R1, J1 and Bruce4, were also male^10-13^. This has limited our understanding of sex-specific pluripotency and impeded the study of female-specific processes, including X chromosome inactivation (XCI); the dosage compensation mechanism in female mammals whereby one of the two X chromosomes becomes stably silenced for the life of the organism^14-17^. In culture, female mESCs appear to be in a more naïve state of pluripotency than male cells, displaying increased expression of naïve pluripotency markers^1, 7, 8^. They also display global hypomethylation compared to males^1-6, 18^, a feature associated with naïve pluripotency. In line with being in a more naïve state, female mESCs are also comparatively slow to exit pluripotency when differentiation is induced^1^. Female mESCs are karyotypically unstable, with XO cells spontaneously arising and then rapidly dominating cultures^3-5, 19^. The development of a defined media for mESC culture has exacerbated and exposed the fragility of female mESCs, likely because this method drives all mESCs towards a more naïve state of pluripotency^20-23^. Known as 2i/LIF^24, 25^, this media provides defined conditions for mESC culture including inhibitors of the glycogen synthase kinase 3β (GSK3β) and MEK/ERK signalling pathways, allowing culture without undefined components such as serum and feeder cells which promote more heterogeneous cell populations. Although challenging for female mESCs, the benefits of a defined media are clear, offering a simpler, more homogenous and reproducible system of mESC culture, while also enabling the study of naïve pluripotency *in vitro*. The use of 2i/LIF media also greatly improves the efficiency of deriving mESC lines *de novo* from pre-implantation embryos^26^, which in turn expands the possibilities for experimental design.

Female mESCs have the unique property of being transcriptionally active from both X chromosomes, a feature shared only with cells of the inner cell mass from which they are derived, induced pluripotent stem cells (iPSC) and primordial germ cells^27-29^. Lineage committed cell types display dosage compensation by XCI. Upon differentiation female mESCs also undergo XCI and following this, hypomethylation and the XO karyotype are no longer features of female cells in culture, with XCI seemingly having a stabilising effect on karyotype and epigenetic makeup^30^. Current evidence suggests that it is the two active X chromosomes that cause female mESCs to behave differently to males, as when female mESCs or iPSCs adopt the XO karyotype, they have similar transcriptomes, epigenomes and differentiation potential to XY pluripotent cells^1, 3, 5, 8, 31^. The mechanisms that lead to these phenotypes are still being uncovered however suppression of the differentiation promoting MAP Kinase pathway is involved^1^, with heterozygous mutation of the X-linked *Dusp9* and *Klhl13* genes able to repress MAP Kinase target gene expression and partly induce the male-like pluripotent state in female cells^2, 7^.

Clearly, the XO karyotype is a preferential state for female mESCs given the domination of this karyotype in cultures, however as it is the XX karyotype that causes female mESCs to behave distinctly to males it is critical that female pluripotency is studied in cultures that maintain a high ratio of XX to XO cells. This is of even higher importance when studying the process of XCI, where undetected XO cells may lead to unreliable conclusions. While the XO issue has been problematic, there have been some seminal studies performed in XX mESCs^1-8, 18^, but there is a large barrier to entry in studying these cells due to a lack of protocols that adopt the latest culture media conditions for mESCs and also enable the retention of both X chromosomes in a high proportion of cells.

Here we exploit the X-linked reporters of our previously published Xmas mESC system^19^, to develop an approach to maximise XX mESCs in fully defined 2i/LIF culture. Further, we find this method to have a stabilising effect also on the karyotype of male mESCs. Our protocol improves the ability to maintain female mESCs *in vitro*, facilitating the study of karyotypically correct cells in a defined media and thus female pluripotency more generally.

## Results

### A method for stabilisation of the XX karyotype in female mESC cultures

We recently reported the Xmas mESC system, which carries X-linked mCherry and GFP reporter constructs driven by the *Hypoxanthine guanine phosphoribosyltransferase (Hprt)* promoter *in trans* to each other, thereby allowing efficient monitoring of XCI by flow cytometry with minimal manipulation of sensitive female mESCs^19^. Moreover, as the X chromosome is biallelically expressed in mESCs, the Xmas reporter alleles may be used to determine the XX/XO karyotype ratio of a given Xmas mESC line. The Xmas reporter alleles are maintained as mouse lines, which when intercrossed produce female offspring with GFP and mCherry marking different X chromosomes when crossed (X^*Hprt*-GFP^ X^*Hprt*-mCherry^, Fig. 1A and S1A). This is important as it allows for the constant rederivation of fresh and karyotypically normal primary Xmas mESC lines. Here we have exploited these properties of the Xmas system to determine an optimised approach to maintaining the XX karyotype in female mESCs. Starting with the current best practice approach for the culture of mESCs in 2i/LIF media^32^, we iteratively and empirically determined which features improved the stability of the XX karyotype through hundreds of rounds of Xmas mESC derivations, to arrive at an optimised protocol when considering XX karyotype. The changes in our method (Keniry2022) compared with the prior published approach (Mulas2019) include increased plating density, increased frequency of passaging (every 24 hours), cells grown in suspension in non-tissue culture treated plates and increased media volumes. We provide the full details of our method as Supplementary Protocol 1.

**Fig. 1.**
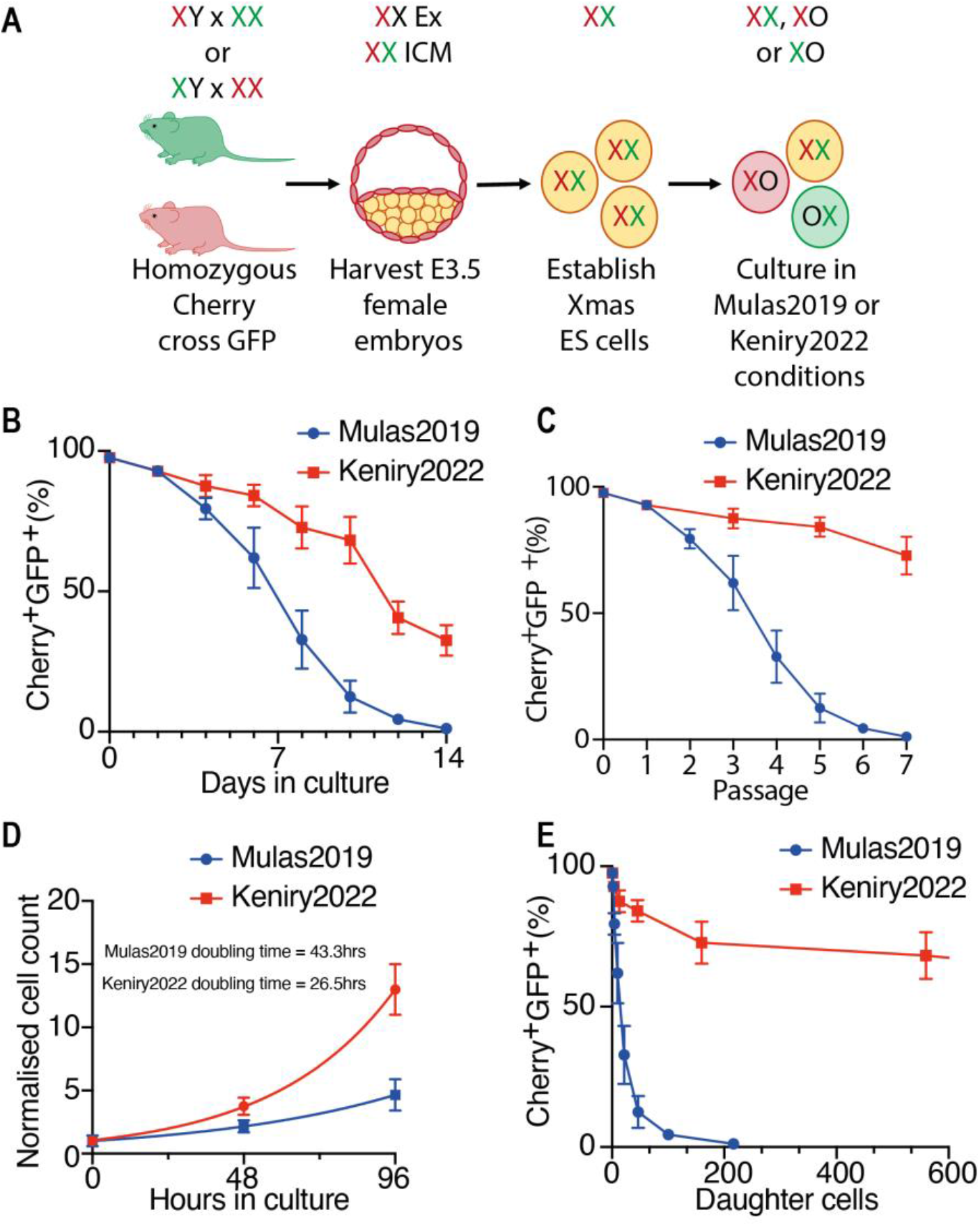
Stabilisation of the XX karyotype in mESC culture. **(A)** Schematic representation of the workflow for using the Xmas mESC system to identify karyotype of female mESCs. **(B)** Flow cytometry data from primary female Xmas mESCs maintained in 2i media for 14 days in either Mulas2019 or Keniry2022 culture conditions, where presence of the reporter alleles indicates the X karyotype of the cell. **(C)** Same data as shown in (B) transformed to reflect passage number instead of days in culture. **(D)** Flow cytometry data indicating cell growth of Xmas mESCs grown under either Mulas2019 or Keniry2022 culture conditions. Lines indicate non-linear fit of data, n=6. Doubling times, calculated from these data, are given. **(E)** Same data as shown in (B) transformed by the calculated doubling time of each method to reflect number of XX daughter cells produced.

Through these subtle changes to the existing best practice method for mESC maintenance we were able to substantially increase retention time of the XX karyotype when compared to the previously published method^24, 32^, either when analysed in terms of days in culture (Figs 1B and S1B) or passage number (Fig. 1C). Moreover, our method also shortens doubling time for XX cells (26.5 hours for Keniry2022 vs 43.3 hours for Mulas2019, Fig. 1D), meaning we can produce substantially more XX daughter cells from each mESC derivation (Fig. 1E). While this method does not completely rescue the XX karyotype, we find that these improvements allow for XX lines to be sufficiently expanded to quantities required for most experimental procedures.

### New method stabilises XX karyotype of F1 mESC lines

We developed our method to stabilise the XX karyotype using Xmas mESCs, which are on the C57/Bl6 strain background. To test whether the method could stabilise karyotype in another strain background, we derived mESC lines from FVB/NJ (FVB)/CAST/EiJ (CAST) F1 blastocysts and passaged these cells for 10 days in either the Keniry2022 or Mulas2019 mESC culture conditions. As these F1 mESC lines lacked the Xmas reporters, we measured X karyotype by DNA florescence *in situ* hybridisation (FISH) with a BAC against the *Huwe1* region of the X chromosome, followed by microscopy and found a substantial stabilisation of the XX karyotype when cells were cultured by our modified method (Fig. 2A,B), suggesting this will improve XX karyotype retention in mESC lines derived from diverse strain backgrounds.

**Fig. 2.**
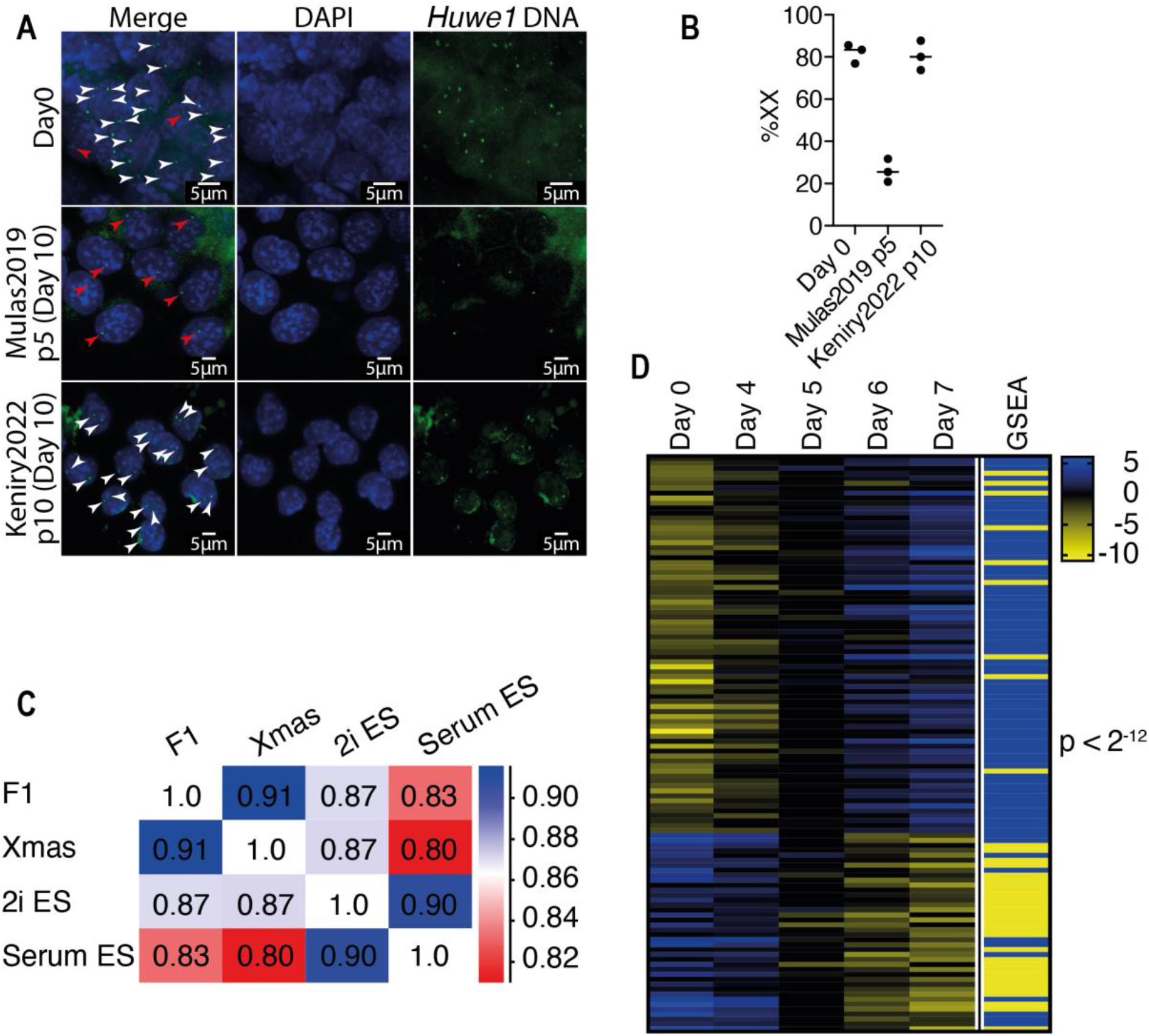
XX karyotype is stabilised in multiple strain backgrounds. **(A)** DNA FISH using a BAC targeted to the *Huwe1* region of the X chromosome in X^FVB^X^CAST^ mESCs cultured in either traditional Mulas2019 conditions for 5 passages (p5, 10 days in culture) or Keniry2022 culture conditions for 10 passages (p10, 10 days in culture). White and red arrows mark the *Huwe1* DNA in cells deemed XX and XO respectively. **(B)** Quantification of data from (A). **(C)** Pearson correlation matrix of our own published RNA-seq from X^FVB^X^CAST^ and Xmas mESCs cultured by our modified conditions together with other publicly available RNA-seq from mESCs grown in serum or mESCs grown in 2i. **(D)** Heatmap of expression values (rpm log_2_) from our published RNA-seq in X^FVB^X^CAST^ mESCs during differentiation, compared to a mESC differentiation gene set (GSEA). Yellow indicates down-regulation, blue up-regulation in GSEA gene set. P-value determined by Chi-square.

### New method does not alter mESC identity

We next sought to identify how the transcriptional status of mESCs may be affected by culture in our conditions, by reanalysing published mESC RNA-seq datasets from Xmas mESCs and FVB x CAST F1 mESCs grown in our conditions, mESCs grown under standard 2i/LIF conditions and mESCs grown in serum-containing media under traditional conditions^19, 20, 33^. All four datasets were highly correlated, however cells grown by our method were more highly correlated with mESCs grown in 2i/LIF than they were to mESCs grown in serum-containing media (Fig. 2C), as expected due to the use of 2i media in our culture conditions. These data show that the naïve pluripotent state expected in 2i mESCs is maintained in our modified method.

We next sought to determine the differentiation potential of mESCs cultured by our modified culture conditions by reanalysis of a published dataset of FVB x CAST F1 mESCs during a differentiation timecourse^19^. We compared these data to a mESC differentiation gene set and found a very high correlation (p<2^-12^, Fig. 2D). These data suggest that mESCs grown under our conditions differentiate with similar transcriptional kinetics to known mESCs. We have previously shown that Xmas mESCs grown under our modified conditions form teratomas containing differentiated cells of all three germ layers upon injection into nude mice^19^.

### Modified culture conditions maintain DNA hypomethylation of XX cells

DNA methylation has a stabilising effect on the genome, therefore we were interested to test whether our modified culture conditions may be maintaining the XX karyotype by relieving the hypomethylation typically observed in XX mESCs. To test this, we grew Xmas mESCs under our culture conditions and separated the XX and XO populations by fluorescence activated cell sorting of the Xmas reporter alleles, then measured DNA methylation by reduced representation bisulfite sequencing (RRBS). This analysis revealed that XX cells remained globally hypomethylated compared to XO cells (Fig. 3A). The pattern of hypomethylation of the XX population was also observed at imprinting control regions and repetitive regions (Line1, SINE and LTR, Fig. 3B). There was no difference in DNA methylation between XX and XO populations at CpG islands, which are already globally hypomethylated compared to the rest of the genome (Fig. 3B). These data suggest that the stabilisation of the XX karyotype achieved by our culture conditions is not due to increased DNA methylation. Therefore, the hypomethylated epigenome of XX female mESCs is maintained by our method, and so these cells are a suitable model in which to study this unique property of female pluripotency. Conversely, our modified method is unlikely to stop the erosion of DNA methylation observed at female imprinting control regions^3, 4^.

**Fig. 3.**
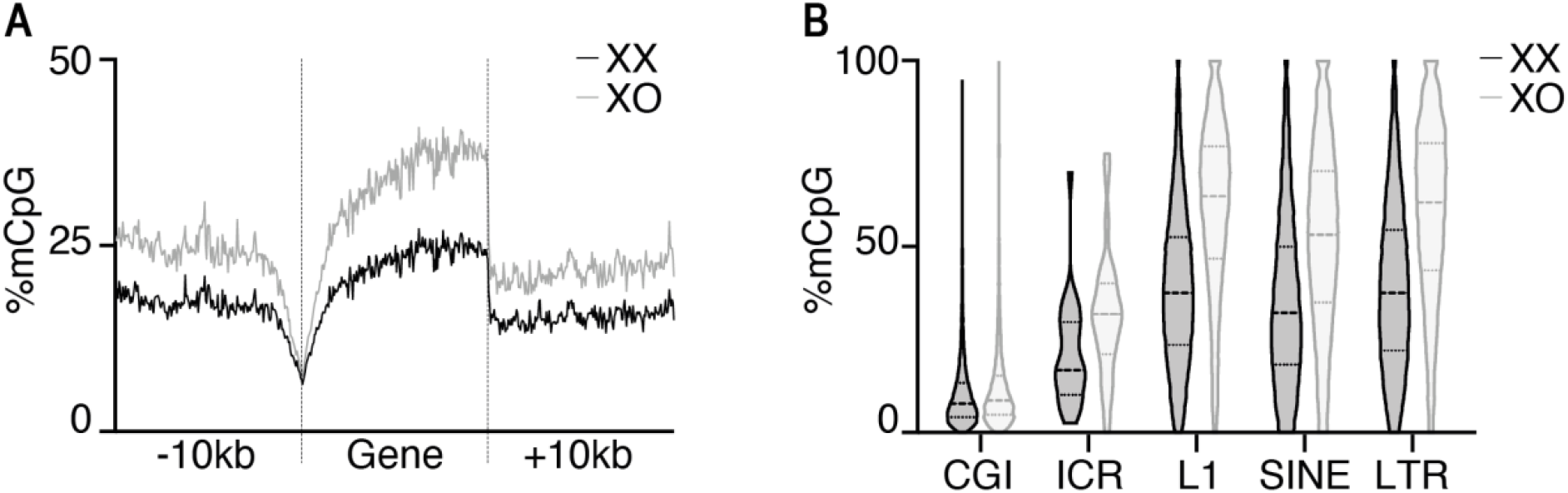
Modified culture conditions maintain female mESC hypomethylation. **(A)** Reduced representation bisulfite data in FACS separated XX and XO Xmas cells cultured by our modified method. The average CpG methylation across all autosomal genes +/-10kb is shown. **(B)** Violin plots of reduced representation bisulfite data in FACS separated XX and XO Xmas cells cultured by our modified method. Bold dotted line indicates median, light dotted lines indicate 25^th^ and 75^th^ centiles. CpG methylation is shown at CpG islands (CGI), imprinting control regions (ICR) and repetitive elements (L1, SINE and LTR).

### Modified culture system improves male mESC fitness

We next tested our culture method on male mESCs, culturing them with our protocol or the current the Mulas2019 mESC protocol^32^, as we were interested to know whether the method could also improve the genetic stability of these more commonly used cells. Note that we present these data passage matched, which most closely aligns the number of cell divisions for each method, however for comparison of days in culture p20 by the Keniry2022 method is analogous to p10 of the Mulas2019 method. Principal component analysis of RNA-seq on these cells showed our protocol maintained male mESCs in a state that is transcriptionally similar to cells freshly derived from the blastocyst, whereas cells under Mulas2019 conditions diverged (Fig. 4A). We identified 5526 differentially expressed genes between the methods, with gene set testing identifying ribosome and mitochondrial associated genes as being significantly upregulated in cells maintained under Keniry2019 conditions (Fig. 4B,C and Table 2), consistent with the more rapid self-renewal that we observe when cells are cultured by our method.

**Fig. 4.**
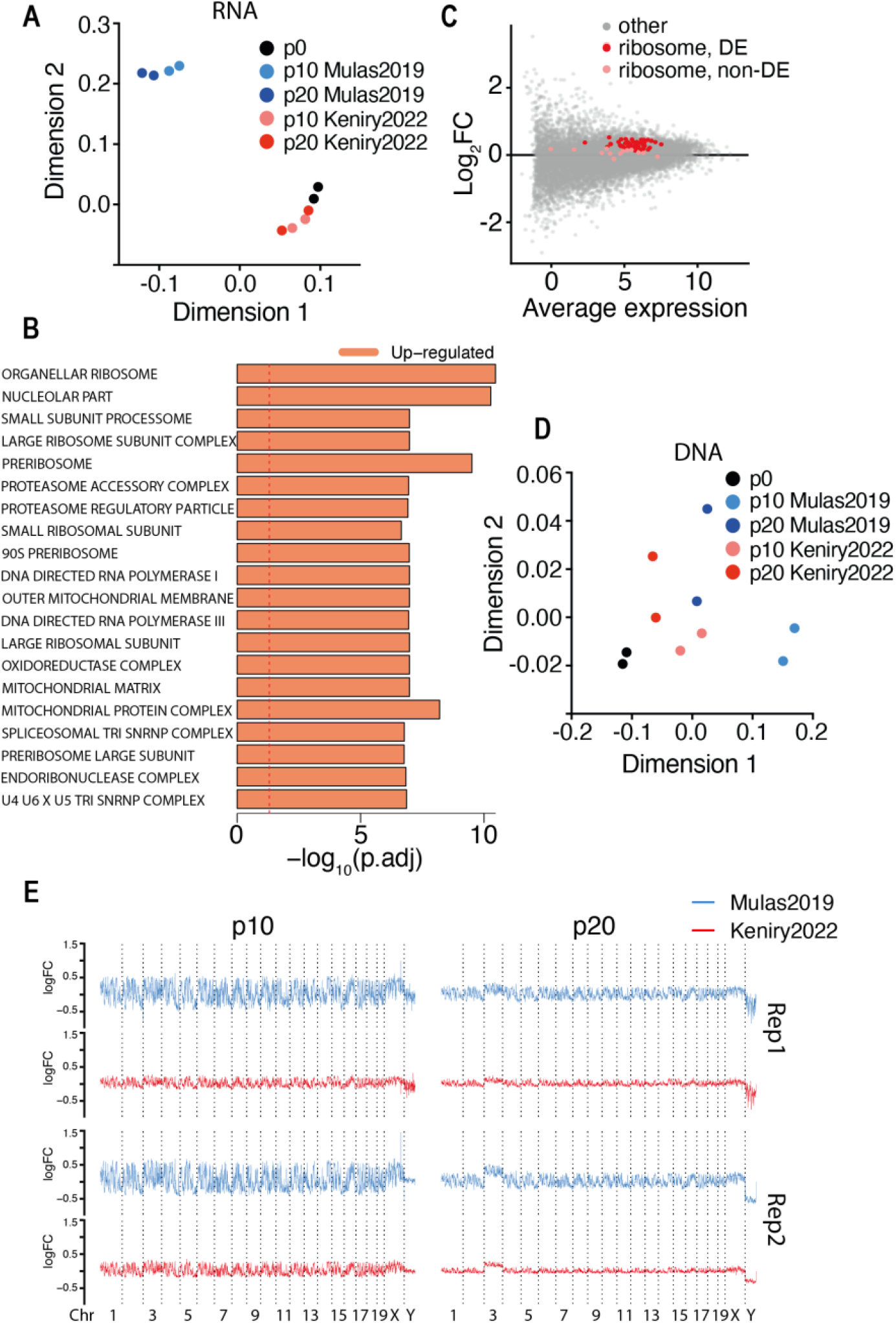
Modified culture conditions maintain transcriptome and karyotype of male mESCs. **(A)** Multi-dimensional scaling (MDS) plot of RNA-seq data from p0 male mESCs compared to cells cultured in either Mulas2019 or Keniry2022 conditions at p10 and p20 (n = 2). Note that the Mulas2019 method has 48h splitting rather than 24h in the Keniry2022 method, thus to compare samples that are days in culture matched, rather than passage matched, p10 Mulas2019 and p20 Keniry2022 are both 20 days in culture beyond p0. **(B)** Results of gene set testing from combined p10 and p20 samples using either the Mulas2019 or Keniry2022 culture conditions. Dashed line indicates p-value = 0.05. **(C)** MA plot showing the average fold change (log_2_FC) at p10 and p20 combined of genes between male mESCs grown in Mulas2019 or Keniry2022 culture conditions. Significantly differentially expressed ribosomal genes are indicated in red and non-significant ribosomal genes in pink. **(D)** MDS plot of DNA-seq data in 1Mb bins from p0 male mESCs compared to cells cultured in either Mulas2019 or Keniry2022 culture conditions at p10 and p20 (n = 2). **(E)** Read coverage plots of 1 Mb bins across all chromosomes of male mESCs cultured in either Mulas2019 or Keniry2022 conditions with reads normalised to equivalent positions in respective p0 samples from two replicate cell lines.

DNA-seq on these cells revealed no major karyotypic abnormalities on a chromosome wide scale (Fig. S2). However, loss of a Y chromosome was observed in one of the cell lines, occurring during the derivation or expansion process, prior to our experimental culture conditions (Fig. S2). We are unsure as to why this might occur, however male Y chromosome instability in cell culture has been reported before^34^. Despite no chromosome-wide differences between the two culture methods, we saw by principal component analysis that autosomes obtained from cells cultured under our conditions remained most similar to that of freshly derived cells during culture, whereas cells cultured with the Mulas2019 method diverged (Fig. 4D); consistent with our method maintaining a more stable karyotype. Analysing the genome in 1Mb bins revealed no differentially represented regions when cells were cultured by our method (FDR < 0.05, log_2_FC > 1.1). By contrast the Mulas2019 method had ∼57% of the genome differentially represented at p10 and ∼5.5% at p20, suggesting prolonged culture selected against major karyotypic abnormalities (Fig. 4E and Table 2). Even when considering equivalent time in culture (p10 Mulas2019 vs p20 Keniry2022), our protocol markedly improves karyotype maintenance. Therefore, our new method improves karyotypic maintenance of both male and female mESCs *in vitro*.

## Discussion

Female and male pluripotency are different, however due to complications with *in vitro* culture female mESCs are severely underrepresented in the literature. Here we have utilised the Xmas mESC system to efficiently detect the presence of X chromosomes and develop a protocol that improves maintenance of the XX karyotype in defined media. Other studies have recently improved culture of female mESCs, achieving significant stabilisation of the epigenome, but were unable to preserve the XX karyotype^2-4^; the feature critical to maintaining the uniqueness of female mESCs and iPSCs^1, 3, 5, 8, 31^. Since our protocol was optimised empirically, we cannot be sure why our conditions better preserve the XX karyotype, but we believe it is due to decreased cell stress, which is an area that could be investigated further in the future. Despite significant improvements, we were unable to completely prevent XO cells eventually becoming predominant in cultures. While our method does not completely preserve the XX karyotype, we find that these improvements allow for XX lines to be sufficiently expanded to quantities required for most experimental procedures. Indeed, we recently utilised our modified mESC culture method, together with Xmas mESCs, to report the first screen for regulators of XCI in the native female context^19^; a feat not previously possible.

Female mESC cultures grown under serum/LIF conditions have been used in XCI studies with success for many years, and although they require regular karyotype monitoring, it seems that on occasion a specific line is able to maintain a more constant XX to XO cell ratio for much longer in serum/LIF than mESCs grown in 2i/LIF media. Cells grown in serum/LIF media are subject to pro-differentiation signals, but activation of JAK-STAT2 signalling by LIF does not allow them to fully differentiate, rather they are maintained in a heterogeneous state of primed pluripotency^35, 36^. It is possible that this heterogenous state relieves the pressure to lose an X chromosome in some cells, leading to an eventual stabilisation of the XX to XO ratio. There are many benefits to using the defined 2i/LIF media to maintain mESCs in a naïve state, including ease of culture, consistency between experiments, homogeneous cell populations and ease of derivation from preimplantation embryos. In some cases, studying the establishment of XCI from the naïve state, as opposed to primed, may be more appropriate depending on the question. It is therefore beneficial to have robust protocols that maintain the XX karyotype of female mESCs in the defined media conditions.

While we were developing our protocol, it was reported that lower concentration of MEK inhibitor partially stabilises the XX karyotype^37^. We have tested this and find low MEK inhibitor further stabilises the Xmas mESC karyotype under our conditions (data not shown), but again does not solve the problem entirely. We note here that another possible strategy for maintaining XX Xmas mESCs is to use FACS sorting to remove XO cells from cultures, however we have found this approach to provide no benefit as the stress of sorting leads to more rapid loss of the XX karyotype once the cells are returned to culture. We suggest a genetic solution to maintaining high quality mESC cultures may be identified by screening in Xmas mESCs for genes whose depletion stabilises the XX karyotype.

Another feature of our culture method is a substantially reduced doubling time for female mESCs from 43.3 hours with the Mulas2019 method to 26.5 hours with the Keniry2022 method. This accelerated doubling time allows for the production of substantially more XX mESCs and over a shorter time period. Reduced cellular stress may also be the contributing factor for the accelerated cell cycle we observe. Previous publications have noted shorter doubling times for mESCs of between 12 and 30 hours depending on culture method^38^. However, these studies, as with the majority of mESC studies, only report findings from male cells. To bring the XX female mESC doubling time to within the range of male mESCs is a significant improvement.

Here we have shown that mESCs cultured by our modified method remain transcriptionally similar to mESCs cultured by the current best practice method. Critically, we also find that our method maintains the DNA hypomethylation that is a signature of XX female mESCs, suggesting that this is a viable approach for study of female pluripotency. In our previous publication using our improved mESC culture protocol we also show that differentiation and XCI proceed as expected and with the epigenetic hallmarks expected of these processes^19^. As XO female mESCs lose the unique properties of female pluripotency, it is critical to maintain high quality cultures of XX female mESCs in defined media for research. Here we provide a protocol that makes this more possible, allowing further discovery of previously hidden facets of this unique female cell type.

## Methods

### Animal strains and husbandry

Animals were housed and treated according to Walter and Eliza Hall Institute (WEHI) Animal Ethics Committee approved protocols (2014.034, 2018.004, 2020.050 and 2020.047). Xmas mice are C57BL/6 background and have been published previously^19^. Castaneus (CAST/EiJ) mice were obtained from Jackson labs and maintained at WEHI. FVB/NJ mice were obtained from stocks held at WEHI. Oligonucleotides used for genotyping are published^19^.

### Derivation of mESCs

Mouse ESCs were derived as previously described^19^. A protocol is supplied as a supplementary method to this manuscript.

### Modified culture method for mESCs (Keniry2022)

We provide a fully detailed lab protocol as a supplementary method to this manuscript. In brief, mESCs were maintained in 2i+LIF medium^24^ [KnockOut DMEM (Life Technologies), 1x Glutamax (Life Technologies), 1x MEM Non-Essential Amino Acids (Life Technologies), 1 X N2 Supplement (Life Technologies), 1 X B27 Supplement (Life Technologies), 1x Beta-mercaptoethanol (Life Technologies), 100 U/mL Penicillin/100 µg/mL Streptomycin (Life Technologies), 10 µg/mL Piperacillin (Sigma-Aldrich), 10 µg/mL Ciprofloxacin (Sigma-Aldrich), 25 µg/mL Fluconazol (Selleckchem), 1000 U/mL ESGRO Leukemia Inhibitory Factor (Merck), 1 µM StemMACS PD0325901 (Miltenyi Biotech), 3 µM StemMACS CHIR99021 (Mitenyi Biotech)] in suspension culture on non-tissue culture treated plates at 37°C in a humidified atmosphere with 5% (v/v) carbon dioxide and 5% (v/v) oxygen. Antibiotic and antimycotics are added to prevent contamination from flushing during the derivation process. Daily passaging of mESCs was performed by allowing unattached colonies to settle in a tube for < 5 minutes. Supernatant containing cellular debris was removed before resuspension in Accutase (Sigma-Aldrich) and dissociation at 37°C for 5 minutes to achieve a single-cell suspension. At least 4 x volumes of mESC wash media [KnockOut DMEM (Life Technologies), 10% KnockOut Serum Replacement (Life Technologies), 100 IU/mL penicillin/100 µg/mL streptomycin (Life Technologies)] were added to the suspension and cells were pelleted by centrifugation at 600 x g for 5 minutes and plated into an appropriately sized non-tissue culture treated plate, never flasks, in an excess of 2i+LIF media. Cells were assessed for XX karyotype regularly by flow cytometry.

We also used a variant of this method where cells were grown on plates coated with 0.1% gelatin, when producing Fig. S1B. Culture conditions were otherwise identical.

### Current best practice culture method for mESCs (Mulas2019)

For comparison of our method we used the current state-of-the-art method for mESC culture^32^, exactly as described apart from the addition of Piperacillin, Ciprofloxacin and Fluconazol to the 2i/LIF media, as described above. Aside from this, all reagents were as for our modified culture method. The primary differences of Mulas2019 culture method are that plating density is decreased, passaging occurs every 48h, cells grow on a gelatin substrate in tissue culture treated plates and media volumes are decreased compared to the Keniry2022 method.

### Differentiation of mESCs

Our mESC differentiation protocol has been published previously^19^. Briefly, we employ undirected differentiation by transitioning cells from 2i+LIF media into DME HiHi media [DMEM, 500 mg/L glucose, 4 mM L-glutamine, 110 mg/L sodium pyruvate, 15% fetal bovine serum, 100 U/mL penicillin, 100 μg/mL streptomycin, 0.1 mM nonessential amino acids and 50 μM β-mercaptoethanol] in 25% increments every 24 hours. Cells continue to differentiate in 100% DME HiHi media. For staging, we refer to the first day of differentiation as being day 0, when cells are placed into 75% 2i/LIF and 25% DME HiHi.

### Flow cytometery analysis and sorting

Cells were prepared in KDS-BSS with 2% (v/v) FBS, with dead cells and doublets excluded by size and analysed using a BD LSRFortessa cell analyser. Cells were prepared similarly for sorting using a FACSAria. Flow cytometry data were analysed using FlowJo.

### RNA-seq library generation and analysis

For the RNA-seq depicted in Fig. 2C,D, we compared published datasets^20, 33^. For the RNA-seq depicted in Fig. 4A-C, we derived male C57/Bl6 mESCs using our culture methods, for two independent lines. These cells were then split in two (p0) and cultured for 10 and 20 passages using either the conditions given in this manuscript or the Mulas2019 method, described above and in^32^. Cells were collected by the addition of lysis buffer and RNA extracted using the Quick-RNA MiniPrep kit (Zymo Research). Next Generation Sequencing libraries were prepared using by TruSeq RNA sample preparation kit (Illumina). Samples were sequenced in-house on the Illumina NextSeq500 platform producing 75bp single-end reads. Quality control and adapter trimming were performed with fastqc and trim_galore^39^ respectively. Reads were aligned to the mm10 reference genome using histat2^40^. Expression values in reads per million (RPM) were determined using the Seqmonk package (www.bioinformatics.babraham.ac.uk/projects/seqmonk/), using the RNA-seq Quantitation Pipeline. Initial data interrogation was performed using Seqmonk.

Gene set testing and differential gene expression analysis was performed by making two groups by pooling samples at all passages from either the Mulas2019 culture method or the Keniry2022 method. Differential expression analysis between the two mESC culture methods was performed on gene-level counts with TMM normalisation, filtering out genes expressed in fewer than half of the samples, using edgeR v3.26.7^41, 42^. Model-fitting was performed with voom v3.40.6^43^ and linear modelling followed by empirical Bayes moderation using default settings. Differential expression results from voom were used for gene set testing with EGSEA v1.12.0^44^ against the c5 Gene Ontology annotation retrieved from MSigDB, aggregating the results of all base methods by ‘fry’ and sorting by median rank.

### DNA-seq library preparation and analysis

We derived male C57/Bl6 mESCs using our culture methods, for two independent lines. Following an 18 day derivation period cells were split in two (p0) and cultured for 10 and 20 passages using either the Keniry2022 conditions given in this manuscript or the Mulas2019 method, described above and in^32^. Note that the Mulas2019 method has 48h splitting rather than 24h in the Keniry2022 method, thus to compare samples that are days in culture matched, rather than passage matched, p10 Mulas2019 and p20 Keniry2022 are both 20 days in culture beyond p0. Sequencing libraries were prepared using the TruSeq DNA sample preparation kit (Illumina) and sequenced in-house on the Illumina NextSeq500 platform with 75bp single-end reads. Reads were mapped to mm10 with bowtie2^45^ and counted in 1Mb bins along the genome using the GenomicAlignments R/Bioconductor package^46^ and computed the percentage of reads mapped to each chromosome. Only bins on the autosomes and sex chromosomes were included and those bins overlapping the ENCODE blacklisted regions were excluded. For each sample, we computed the coverage of each bin in log counts per million. We then computed the log fold changes comparing each sample to the relevant p0 sample and plotted these by bin position along the genome. We used the edgeR R/Bioconductor package^41^ to perform a multidimensional scaling plot of distances between samples based on the log fold changes. Differential abundance analysis was performed using edgeR and limma^47^. Briefly, the voom method^43^ was used to prepare count data for linear modelling and the within-cell line correlation estimated using the ‘duplicateCorrelation’ function from the limma package^48^. The voom method was then re-applied (now accounting for the within-cell-line correlation), the within-cell-line correlation re-estimated, and these transformed data used as input to a linear model with design matrix encoding the passage number and protocol of each sample while blocking on the cell line and including the estimated within-cell-line correlation when fitting the linear models. We used the empirical Bayes statistics^49^ to test for differential abundance at p10 vs. p0 and p20 vs. p0 within each protocol at a false discovery rate of 0.05 and requiring a minimum log2-fold change of 1.1^50^.

### DNA fluorescence *in situ* hybridisation (FISH)

DNA FISH was performed as previously described^51^ on female mESCs derived by crossing FVB/NJ (FVB) dams with CAST/EiJ (CAST) sires, then passaged either with Keniry2022 or Mulas2019 methods. The X chromosome was detected with a BAC probe against the *Huwe1* region (RP24-157H12), as previously described^52^. The probe was labelled with Green-dUTP (02N32-050, Abbott) by nick translation (07J00-001, Abbott). The cells were mounted in Vectashield antifade mounting medium (Vector Laboratories) and visualised on LSM 880 or LSM 980 microscopes (Zeiss). Images were analysed using the open source software FIJI^53^.

### Reduced representation bisulfite sequencing

Xmas mESCs were cultured for 7 passages using our modified method, before being FACS separated into XX and XO populations based on the Xmas reporter alleles, using a BD FACSAria III Cell Sorter. Reduced representation bisulfite libraries were prepared and analysed as previously described^54^.

### Accession Numbers

All next generation sequencing data generated for this project have been deposited in the Gene Expression Omnibus (GEO) database under accession number GSE162965.

## Supporting information

Supplementary Protocol

## Acknowledgements

This study was supported by an Australian Research Training Program scholarship (NJ), the Bellberry-Viertel Senior Medical Research Fellowship (MEB), and an Australian National Health and Medical Research Council fellowship (MER). Grant support was provided by the Australian National Health and Medical Research Council (1059624 to MEB, 1140976 to MEB, MER and AK, to MEB 1194345), the Dyson Bequest and the DHB Foundation. Additional support was provided by the Victorian State Government Operational Infrastructure Support, Australian National Health and Medical Research Council IRIISS grant (9000433).

## Contributions

AK and MEB conceived the study. AK, NJ, KAB, MI, and TB performed experiments. AK and PFH performed bioinformatic analysis. AK, MER and MEB secured funding. MER and MEB supervised the project. AK and MEB wrote the manuscript with contribution from all authors.

## Declaration

The authors declare no competing interests.

---

**Fig. S1.**
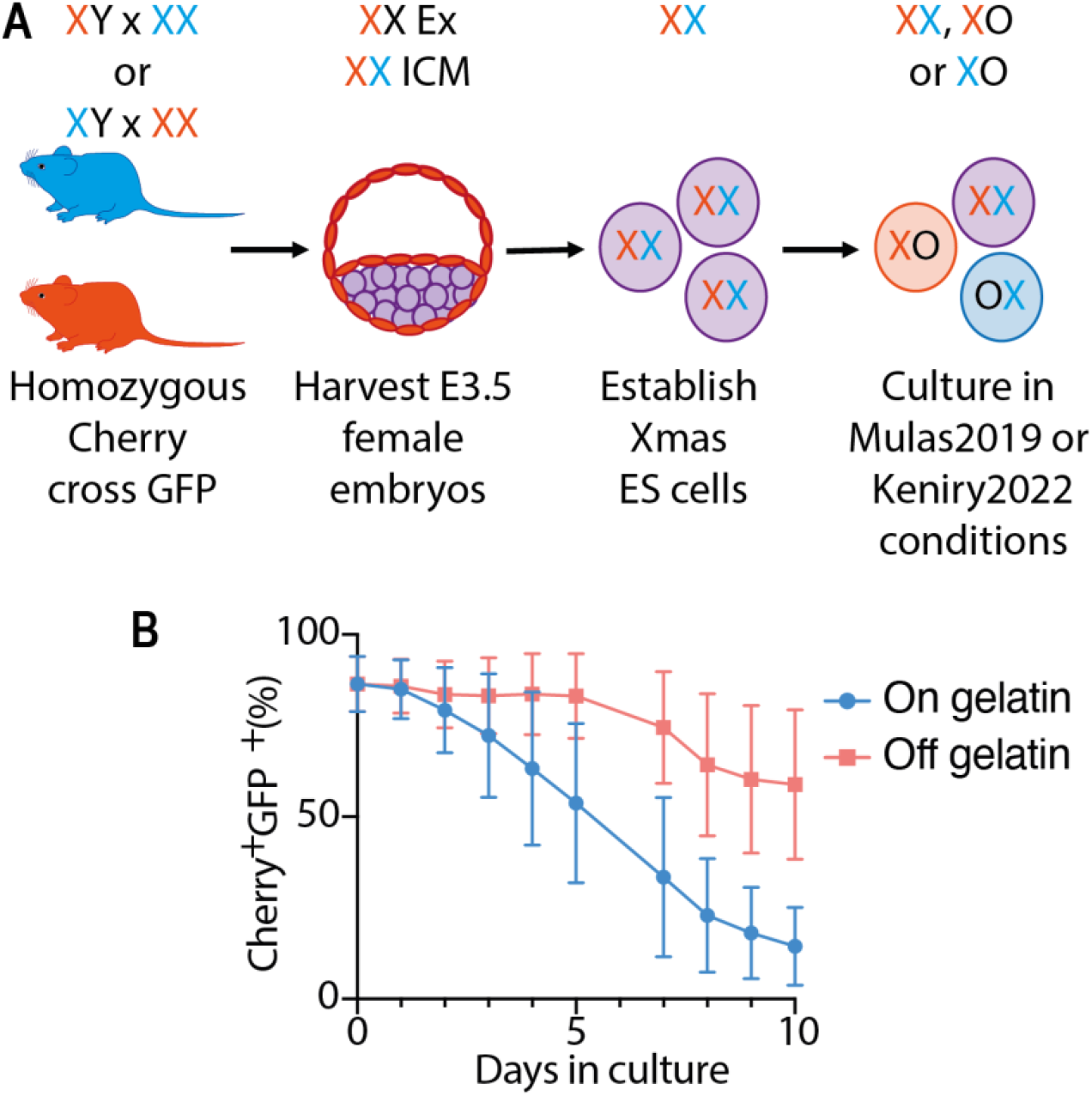
Development of modified culture conditions. **(A)** Colour blind friendly version of Fig. 1b. mCherry is represented in orange and GFP in blue. **(B)** Flow cytometry data from primary female Xmas mESCs maintained in 2i media for 10 days either attached to gelatin (On) or without an attachment substrate (Off), both passaged daily, where presence of the reporter alleles indicates the X karyotype of the cell.

**Fig. S2.**
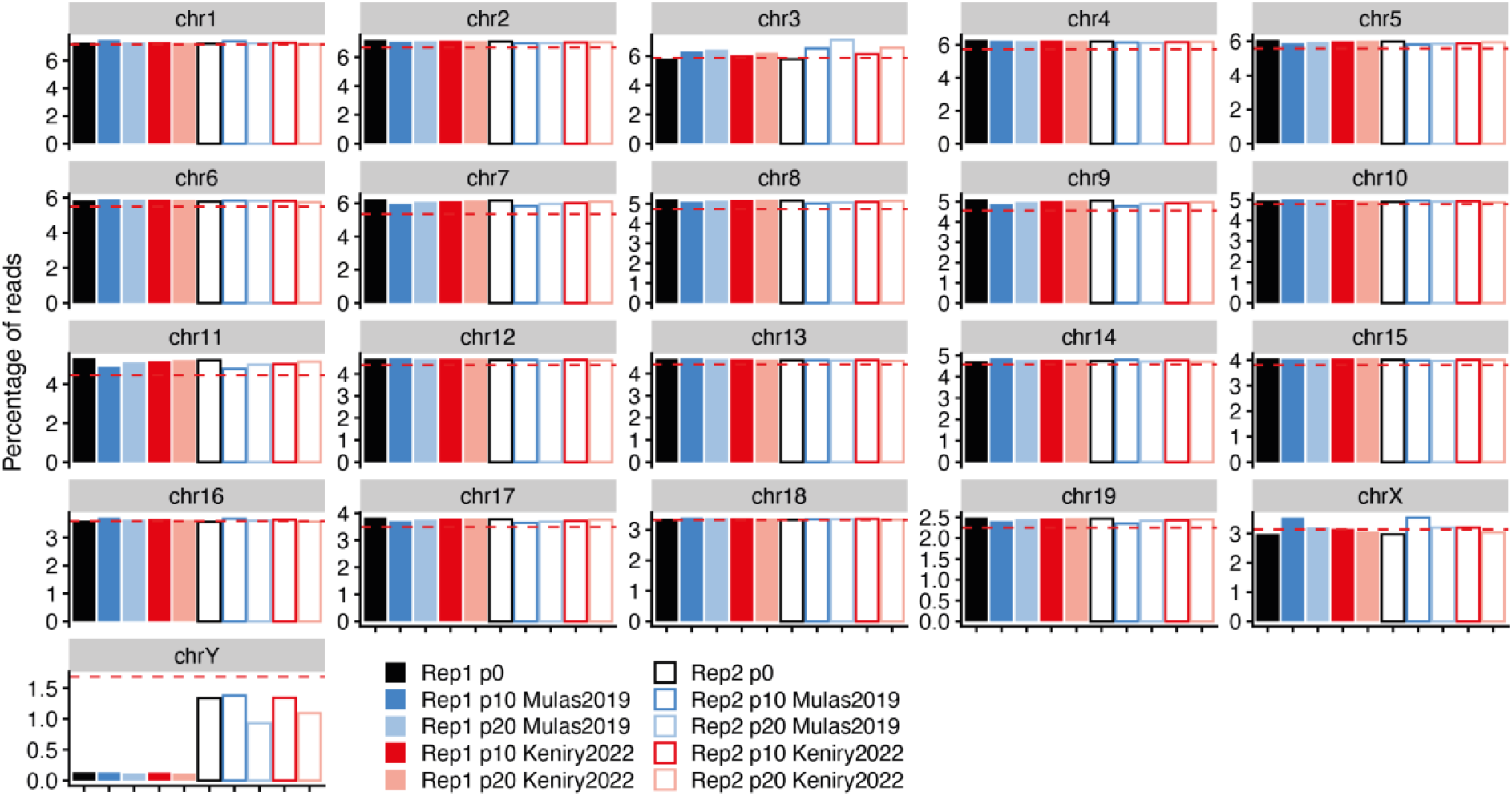
No chromosome wide abnormalities in male mESCs cultured by Mulas2019 or Keniry2022 conditions. Bar graphs showing DNA-seq data from male mESCs maintained in 2i media with the percentage of total reads mapping to the indicated chromosomes at p0 or following p10 or p20 passages in either Mulas2019 or Keniry2022 culture conditions. Dotted lines indicate the expected percentage of reads that should map to the chromosome based on chromosome length. Data for two independent male mESC lines are shown. Note that the Mulas2019 method has 48h splitting rather than 24h in the Keniry2022 method, thus to compare samples that are days in culture matched, rather than passage matched, p10 Mulas2019 and p20 Keniry2022 are both 20 days in culture beyond p0.

**Table. 1 Differential gene expression analysis of RNA-seq data from male mESC**

Table provides the results of differential gene expression between male mESCs cultured with either Mulas2019 or Keniry2022 conditions. p10 and p20 samples have been considered replicates in this analysis.

**Table. 2 Differential genomic representation analysis of DNA-seq data from male mESC**

Table provides the results of differential genomic representation of 1Mb bins between male mESCs cultured with either Mulas2019 or Keniry2022 conditions. The logFC value and the adj.P.Val compares the indicated sample with the equivalent p0 sample.

**Supplementary Protocol 1. Modified mESC Culture Protocol**

This note contains a full lab working protocol for the Keniry2022 culture method of mESCs.

## Notes

### Competing Interest Statement

The authors have declared no competing interest.

