## Supplementary Protocol for "A method for stabilising the XX karyotype in female mESC cultures"

### Xmas mESC methods

#### Materials

- Superovulated pregnant females at E3.5 following $X^{\mathit{Hprt}\text{-mCherry}}X^{\mathit{Hprt}\text{-mCherry}}$ x $X^{\mathit{Hprt}\text{-GFP}}Y$ or $X^{\mathit{Hprt}\text{-GFP}}X^{\mathit{Hprt}\text{-GFP}}$ x $X^{\mathit{Hprt}\text{-mCherry}}Y$ inter-crosses.
- Benchtop centrifuge
- Incubator set to 37°C, 5% $O_2$, 5% $CO_2$ and humidified.
- Dissection microscope
- Non tissue culture treated plates (24-well, 12-well and 6-well).
- 96-well round bottom plate
- IVF petri dishes
- Mouth pipette
- Dissecting tools
- 25-gauge needles
- Xmas wash media (see section **Xmas wash media**)
- 2i Media (see section **2i Media**)
- M2 media
- Phosphate buffered saline (PBS)
- Trypsin-EDTA
- Accutase™

#### Notes before starting

- Best results are achieved when these C57BL/6 strain females receive folligon for superovulation between the ages of 28 and 32 days. This changes with different strain backgrounds.
- The direction of the inter-cross is not important and will yield similar Xmas mESCs regardless of which strain provides the dam and which provides the sire.
- It is critical to always work quickly to minimise the time cells are out of the incubator.
- Utilise aseptic technique as much as possible, however we find it more comfortable to perform microscopy and mouth pipetting outside of a tissue culture hood, but with freshly cleaned equipment.
- Note that we use antibiotic and antimycotic in our 2i media as unwanted yeast and bacteria are frequently obtained when flushing uterine horns. We have tested multiple antibiotics and antimycotics and while the ones listed below best maintain the XX karyotype, better maintenance is achieved when these are left out of the media.
- Once genotyped, individual Xmas mESC outgrowths can be pooled to increase cell numbers more rapidly.
- Limit the amount of times an incubator is accessed while culturing Xmas mESCs as fluctuations in temperature and atmosphere (low oxygen) will negatively impact results. Ideally only open the incubator during one period per day.
- Cells more stably maintain the XX karyotype when cultured at a higher density, compared to a lower density. Err on the side of higher density when deciding how to split cells.
- Keep passaging times as close to 24 hours apart as possible. Less than 24 hours results in a low density of cells at passaging, while longer leads to yellowing of the media and clumping of the cells, both of which promote loss of the XX

karyotype.

- Never pre-warm the bottle of Accutase as it loses potency rapidly when out of the fridge. Instead, aliquot and pre-warm to 37°C.
- Pre-aliquot and warm only the required amount of 2i media to 37°C.
- Xmas mESCs must be cultured in round wells of non-tissue culture treated plates, as the round wells encourage cells to cluster in the middle of the well. Cells cultured in square flasks will disperse throughout the flask, promoting loss of the XX karyotype as local confluency is decreased.
- Perform desired experimentation as soon as the required number of cells is required. If a cell line drops below 80% XX we discard the line and do not experiment on it.
- We use this protocol for deriving and culturing all mESC lines, not just Xmas mESC lines, with the exception that lines which do not carry the Xmas reporter alleles must be sexed and genotyped by PCR.

**Harvesting blastocysts**

1. Add 500 µL of 2i medium to each well of a non-tissue culture treated 24-well plate, allowing one well for each expected blastocyst. For each pregnant female, add 1 mL of 2i media to two 3.5cm IVF petri dishes and 1mL M2 medium to 2 IVF petri dishes. Allow media to equilibrate in 37°C, 5% $CO_2$, 5% $O_2$ humidified incubator for at least 30 minutes.
2. Euthanise pregnant females and remove uterine horns using sterile technique. Store uterine horns in room temperature PBS until all dissections are complete.
3. Using the equilibrated M2 media, flush blastocysts from uterine horns using a 25-gauge needle back into the IVF petri dish. Use a new petri dish for each pregnant female.
4. Under a dissection microscope, wash blastocysts by sequentially mouth-pipetting embryos into a clean pre-prepared and equilibrated petri dish of M2 media, then to a petri dish of 2i media and finally into another petri dish of 2i media. Take care not to transfer debris.
5. Plate embryos into individual wells of the 24-well plate with equilibrated 2i media and incubate in 37°C, 5% $CO_2$, 5% $O_2$ humidified incubator for 7 days.

**Dissociating outgrowths**

1. After 7 days a prominent inner cell mass outgrowth should be visible. For each outgrowth, add 500 µL of 2i media to a well of a non-tissue culture treated 24-well plate, and equilibrate in 37°C, 5% O2, 5% CO2 humidified incubator.
2. For each outgrowth, in a 96-well round bottom plate prepare a well of PBS, followed by a well of Trypsin, a well of Xmas Wash media and a well of 2i media. Use a volume of 100 µL for each media. For best results, this plate should remain at room temperature.
3. Pick outgrowths by mouth pipetting under a dissection microscope and transfer sequentially through PBS, Trypsin, Xmas Wash media and 2i in the pre-prepared 96-well plate. The outgrowths will need 5-8 minutes in the Trypsin well at room temperature and must be monitored under the microscope. Once there are signs that the outgrowth is beginning to dissociate it can be moved to the Xmas Wash well, then immediately to the well containing 2i media.
4. Once in 2i media, use a P200 pipette to dissociate the outgrowth into small clumps of cells by pipetting up and down 2-5 times before transferring media containing cells into the equilibrated 24-well plate of 2i media. Note that at this

point a small portion of single cells from each outgrowth can be used for sex genotyping by flow cytometry. This is our preferred method as it allows us to pool multiple female outgrowths, thereby hastening the time taken to grow the required number of cells for an experiment.

5. Incubate in 37°C, 5% $CO_2$, 5% $O_2$ humidified incubator. Xmas mESC lines are now established and should be passaged as below.

**Maintenance of Xmas mESCs (Improved mESC protocol)**

1. Passage Xmas mESCs every 24 hours in culture (note that the first passage following derivation is often better performed following 48 to 72 hours post derivation).
2. Pre-warm appropriate amount of 2i, Accutase and Xmas Wash media (see section **Media volumes for maintaining Xmas mESCs**).
3. Pipette Xmas mESCs and media out of their well and into a 10 ml centrifuge tube. Centrifuge at 500 rpm for 1 minute. Remove and discard all but the last ~50 µl of media.
4. Add Accutase (500 µl for a small well up to 2 ml for a full 6-well plate), resuspend cells by swirling (not pipetting) and incubate at 37°C for 3-5 minutes.
5. Using a 1 ml pipette triturate cells 6 times to produce a single cell suspension and confirm under a microscope.
6. Add 3 volumes of warm Xmas Wash media and invert to mix.
7. Spin at 1,500 rpm for 3 minutes and remove supernatant by tipping. Note, when passaging low numbers of cells the last drop should be blotted with a tissue to avoid significantly diluting the 2i media.
8. Resuspend cells in 2i media (see section **Media volumes for maintaining Xmas mESCs**) and transfer to an appropriate size well of a non-tissue culture treated plate (see section **Typical passaging scheme**).
9. Incubate in 37°C, 5% $CO_2$, 5% $O_2$ humidified incubator.
10. Passage cells every 24 hours.

**Media volumes for maintaining Xmas mESCs**

Culture Xmas mESCs in an abundance of 2i media, as media that has yellowed promotes loss of the XX karyotype. Typical media volumes for each well size are below:

| | |
|---|---|
| 24-well | 1 ml |
| 12-well | 2 ml |
| 6-well | 5 ml |

**Typical passaging scheme**

Xmas mESCs must be cultured in round wells of non-tissue culture treated plates. Xmas mESCs double approximately every 24 hours, so typically cells are passaged into twice the number of wells they began in, however if their density will not reach $2x10^5$ cells/ml, then they should be returned to a smaller sized well. Note, that cells should not be counted during passaging as the time taken to do this will promote loss of the XX karyotype. Instead, it is necessary to learn to judge cell density by eye. Successful Xmas mESC derivations tend to follow a similar routine of passaging following the dissociation of outgrowths, as follows:

| | |
|---|---|
| Day 1 | 10 - 40 female Xmas outgrowths dissociated and genotyped, and all females pooled into 1x well of a 24-well plate. |
| Day 2 | No passage required |
| Day 3 | Passage back into 1x 24-well |
| Day 4 | Passage into 2x 24-wells |
| Day 5 | Passage into 1x 12-well |
| Day 6 | Passage into 2x 12-wells |
| Day 7 | Passage into 1x 6-well |
| Day 8 | For each additional day double the number of wells of a 6-well plate |

This scheme is intended as a guide and should be adapted to each Xmas mESC derivation.

**Inhibitors and drugs**

- **MEKi (PD0325901, 10mM)** Add 415 µl of DMSO to 2 mg vial of MEKi. Invert to mix and aliquot into 25 µl aliquots. Store at -20°C.
- **GSKi (CHIR99021, 10mM)** Add 430 µl of DMSO to 2 mg vial of GSKi. Warm to 37°C for 5 mins. Invert to mix and aliquot into 150 µl aliquots. Store at -20°C.
- **Fluconazole (5 mg/ml)** Add 1 ml of DMSO to 5 mg powdered fluconazole and mix well. Aliquot into 50 µl aliquots and store -20°C.
- **Ciprofloxacin/Piperacillin (10 mg/ml)** Dissolve 50 mg of powdered Ciprofloxacin and 50 mg of powdered Piperacillin in 5 ml of water. Aliquot into 500 µl aliquots and store -20°C.

**2i media (for 500 ml)**

| | |
|---|---|
| 465 ml | Knockout DMEM |
| 10 ml | Glutamax (100x) |
| 5 ml | Pen/Strep (100x) |
| 5 ml | Non-essential Amino Acids (100x) |
| 900 µl | 2-mercaptoethanol (55 mM) |
| 5 ml | N2 supplement (100x) |
| 10 ml | B27 supplement (50x) |
| 500 µl | Piperacillin/Ciprofloxacin (10 mg/ml each) |
| 1 µl | Fluconazole (5 mg/ml) |
| 25 µl | MEKi (10 mM) |
| 150 µl | GSKi (10 mM) |
| 50 µl | ESGRO/LIF ($10^7$ units/ml) |

- Filter sterilise, wrap in foil and store at 4°C for up to a month.

**Xmas Wash (for 500ml)**

| | |
|---|---|
| 450 ml | Knockout DMEM |
| 50 ml | Knockout serum replacement |
| 5 ml | Pen/Strep (100x) |

- Filter sterilise and freeze at -20°C in 50 ml aliquots.
